# SMCR8 negatively regulates AKT and MTORC1 signaling to modulate lysosome biogenesis and tissue homeostasis

**DOI:** 10.1101/485243

**Authors:** Yungang Lan, Peter M. Sullivan, Fenghua Hu

**Affiliations:** Department of Molecular Biology and Genetics, Weill Institute for Cell and Molecular Biology, Cornell University, Ithaca, NY 14853, USA; Key Laboratory of Zoonosis Research, Ministry of Education, College of Veterinary Medicine, Jilin University, Changchun, China

**Author notes:** To whom correspondence should be addressed: Fenghua Hu, 345 Weill Hall, Ithaca, NY 14853.

**Keywords:** AKT, amyotrophic lateral sclerosis, autophagy, C9orf72, frontotemporal lobar degeneration, inflammation, lysosome, MTORC1, SMCR8, WDR41

## Abstract

The intronic hexanucleotide expansion in the *C9orf72* gene is one of the leading causes of frontotemporal lobar degeneration (FTLD) and amyotrophic lateral sclerosis (ALS), 2 devastating neurodegenerative diseases. C9orf72 forms a heterodimer with SMCR8 (Smith-Magenis syndrome chromosome region, candidate 8) protein. However, the physiological function of SMCR8 remains to be characterized. Here we report that ablation of SMCR8 in mice results in splenomegaly with autoimmune phenotypes similar to that of C9orf72 deficiency. Furthermore, SMCR8 loss leads to a drastic decrease of C9orf72 protein levels. Many proteins involved in the macroautophagy-lysosome pathways are downregulated upon SMCR8 loss due to elevated activation of MTORC1 and AKT, which also leads to increased spine density in the *Smcr8* knockout neurons. In summary, our studies demonstrate a key role of SMCR8 in regulating MTORC1 and AKT signaling and tissue homeostasis.

## Introduction

Hexanucleotide repeat expansion in the *C9orf72* gene is a prevalent genetic cause for frontotemporal lobar degeneration (FTLD) and amyotrophic lateral sclerosis (ALS), 2 devastating neurodegenerative diseases [1, 2, 3]. Reduced expression of the *C9orf72* gene is proposed to be one of the disease mechanisms [4, 5, 6]. However, the cellular function of *C9orf72* remains elusive. Recently, we and others have found that C9orf72 protein forms a complex with 2 additional proteins of unknown functions, SMCR8 and WDR41 [[7, 8, 9, 10, 11, 12]. An early characterization of C9orf72 and SMCR8 by structural prediction suggested that they both contain DENN (<u>d</u>ifferentially <u>e</u>xpressed in <u>n</u>ormal and <u>n</u>eoplastic) domains, which are commonly found in RAB GTPase <u>g</u>uanine nucleotide <u>e</u>xchange <u>f</u>actors (GEFs) [13, 14]. Several RABs have been identified to be the target of C9orf72-SMCR8, including RAB5, RAB7, RAB7L1, RAB8, RAB11 and RAB39 [7, 10, 15, 16, 17]. Additionally, C9orf72 and SMCR8 have been shown to regulate various aspects of the autophagy pathway, despite inconsistent results between different studies [7, 8, 9, 10, 11, 12. These data support the idea that the C9orf72 complex is an important regulator of membrane trafficking. Ablation of C9orf72 in mice results in severe inflammation and autoimmunity [8, 18, 19, 20]. However, the in vivo function of SMCR8 is still unclear.

To study the C9orf72 complex in more mechanistic detail and to investigate the physiological functions of SMCR8, we generated *smcr8* knockout mice in addition to our *c9orf72* knockout mice previously characterized [8]. We found that SMCR8 deficiency in mice causes abnormal inflammatory phenotypes and autoimmunity similar to that of C9orf72 deficiency. Moreover, loss of SMCR8 enhances MTORC1 and AKT activities, decreases lysosomal biogenesis and increases spine density, suggesting that SMCR8 negatively regulates AKT-MTORC1 signaling to maintain tissue homeostasis.

## Results

### Generation of SMCR8-deficient mice

To study the in vivo functions of *Smcr8*, we created *smcr8* knockout mice using the CRISPR-Cas9 system [21, 22]. A mouse line with a 128-base pair (bp) deletion just after the start codon of *Smcr8*, resulting in a frame shift and early stop codon after 22 amino acids was used in our study (Fig. 1A). SMCR8-deficient mice do not have any apparent growth defects (data not shown) but display an obvious spleen enlargement similar to that of C9orf72 deficiency [18, 20]. Splenomegaly starts to appear in the 2-month-old SMCR8-deficient mice, and this phenotype becomes apparent at 4 months of age, with mice having a spleen 2-3 times the size of the littermate control (Fig. 1B).

**Figure 1.**
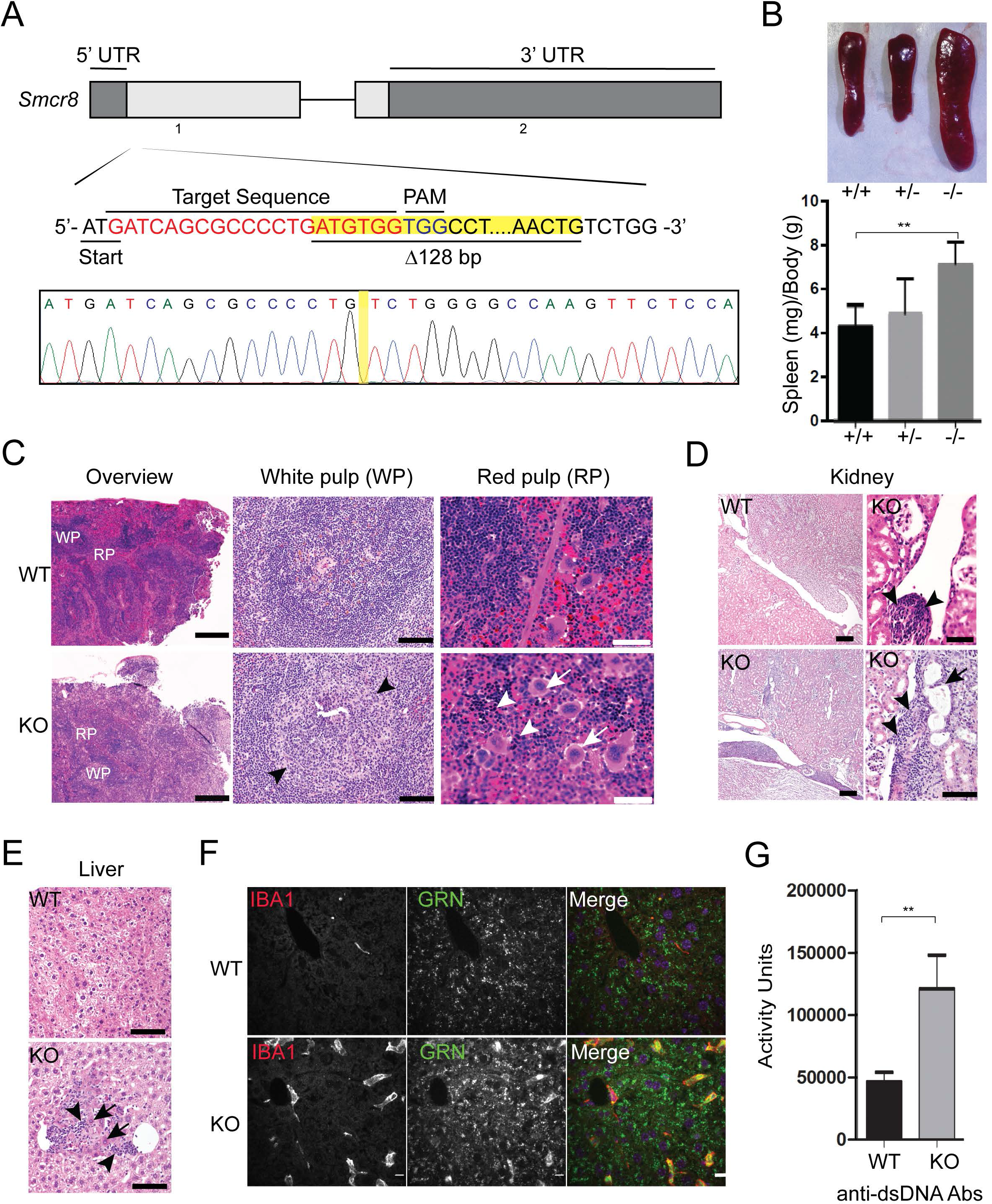
SMCR8 deficiency in mice results in splenomegaly and autoimmunity. (**A**) Schematic drawing of the mouse *Smcr8* gene exon 1 and 2, and the site targeted for editing by CRISPR-Cas9. Sequencing traces of the edited *Smcr8* gene from genomic PCR show 128-bp deletion (highlighted with yellow) near the Cas9 cleavage site. (**B**) Representative images of spleens and quantification of spleen weight from 4 month-old WT and *smcr8*^*−/−*^ mice. n= 3, **: p<0.01, student’s t-test. (**C**) H&E staining of spleen tissues from 12 month-old WT or *smcr8*^*−/−*^ mouse with higher magnification images of white pulp (WP) and red pulp (RP). WP, arrowheads indicate expanded germinal centers in the KO spleen; RP, arrows indicate megakaryocytes and arrowheads indicate erythroid precursors. Scale bar: 500 μm (100 μm and 50 μm in the zoomed in images for WP and RP, respectively). (**D**) H&E staining of kidney tissue from 12-month-old mice. Lymphocytes and macrophage infiltrates (arrowheads) are detected in the interstitium around the pelvis and multifocally in the cortex and medulla in the *smcr8*^*−/−*^(KO) kidney. A dilated tubule (arrow) and other tubules with slightly basophilic cytoplasm are also observed in the *smcr8*^*−/−*^ kidney. Scale bar: 100 μm (50 μm in the zoomed in images for KO kidney) (**E**) H&E staining of liver from 12-month-old mice showing infiltrates of lymphocytes and macrophages (arrowheads) in the *smcr8*^*−/−*^ liver. Arrows indicate hypereosinophilic hepatocytes that might be undergoing degeneration and necrosis. Scale bar: 20 μm. (**F**) Immunostaining of 12-month-old liver sections of WT and *smcr8*^*−/−*^ mice with anti-IBA1 and GRN (granulin) antibodies. Scale bar: 10 μm. (**G**) ELISA to measure anti-dsDNA antibodies (Abs) in serum obtained from 4-months-old WT and *smcr8*^*−/−*^ (KO) mice. n= 3−7, **: p<0.01, student’s t-test.

Hematoxylin and eosin (H&E) staining of the SMCR8-deficient spleen reveals increased white pup to red pulp ratios (∼4:1 compared to ∼1.5:1 in WT) (Fig. 1C). High magnification shows expanded peri-arteriolar lymphoid sheaths in the germinal center (GC) (Fig. 1C). Infiltrated lymphocytes and macrophages were observed in *smcr8*^*−/−*^ liver and kidney (Fig. 1D and E, arrowheads). The presence of macrophages in the liver is confirmed by anti-IBA1 staining (Fig. 1F). The surrounding hepatocytes are hypereosinophilic and sometimes shrunken, potentially undergoing degeneration and necrosis (Fig. 1E). The *smcr8*^*−/−*^ kidney has multifocal perivascular aggregates of lymphocytes and macrophages with similar cells expanding the peri-pelvic interstitium (Fig. 1D). Focally a radiating band of cortex is also expanded by lymphocytes and macrophages with occasionally dilated tubules and tubules with slightly basophilic cytoplasm (Fig. 1D). Significant accumulation of anti-dsDNA antibodies were detected in SMCR8-deficient mice compared to littermate WT controls (Fig. 1G), suggesting that SMCR8 deficiency also results in autoimmunity similar to that of C9orf72 deficiency. Despite inflammation in multiple peripheral organs, microglia do not show any obvious changes in number and morphology in the SMCR8-deficient mice (Fig. S1).

### SMCR8 and C9orf72 stabilize each other

Because C9orf72, SMCR8, and WDR41 form a complex, we hypothesize that they might stabilize each other. We found that loss of SMCR8 (Fig. 2A and B, S2) and to a lesser extent, loss of WDR41 (Figure 2F, 2G) led to a significant decrease in C9orf72 levels in brain lysates and fibroblasts. mRNA analysis did not show any changes in *C9orf72* mRNA levels in *smcr8*^*−/−*^ samples (Fig. 2C), suggesting the reduction of C9orf72 protein levels in *smcr8*^*−/−*^ mice is likely due to post-transcriptional mechanisms. Furthermore, SMCR8 protein levels were also dramatically decreased in brain lysates and fibroblasts derived from *c9orf72*^*−/−*^ and, to a lesser extent, *wdr41*^*−/−*^ mice (Fig. 2D-2G). These data are consistent with our previous finding that C9orf72 forms a heterodimer with SMCR8, which subsequently interacts with WDR41 [8]. Unfortunately, we cannot assess WDR41 level changes due to lack of available specific antibodies.

**Figure 2.**
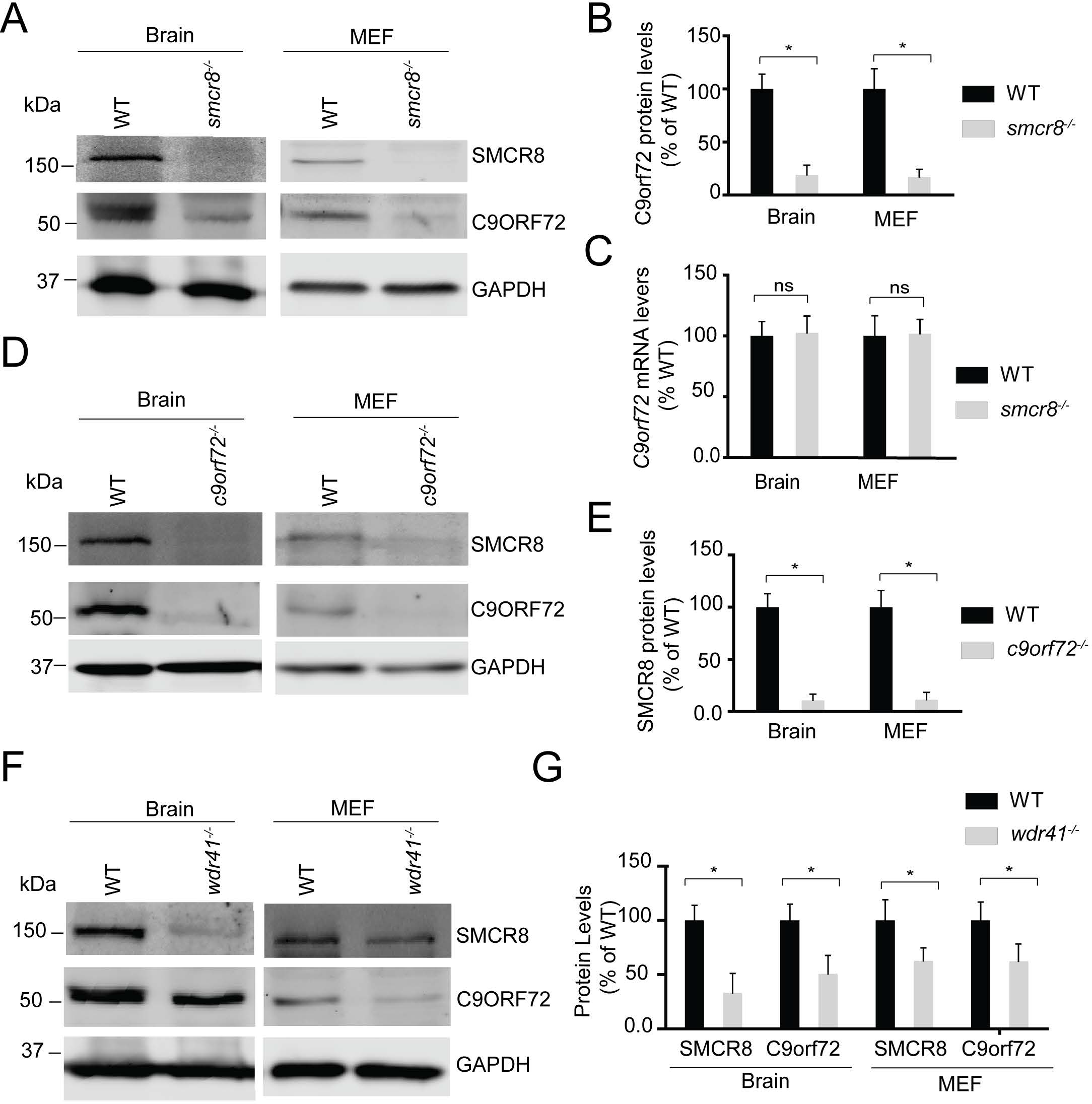
C9orf72 and SMCR8 require each other for stability, and the complex is stabilized byWDR41. (**A**) Western blot analysis of SMCR8 and C9orf72 proteins in WT and *smcr8*^*−/−*^ lysates as indicated. (**B**) Quantification of C9orf72 protein levels for the experiment in (A). C9orf72 levels were quantified and normalized to GAPDH (n=3, *, p<0.05, student’s t-test). (**C**) qPCR analysis of *c9orf72* mRNA levels in WT and *smcr8*^*−/−*^ brain and MEF samples. (**D**) Western blot analysis of C9orf72 and SMCR8 protein in WT and *c9orf72*^*−/−*^ brain and MEF cell lysates. (**E**) SMCR8 levels were quantified and normalized to GAPDH for experiment in (D) (n=3, *, p<0.05, student’s t-test). (**F**) Western blot analysis of C9orf72 and SMCR8 proteins in WT and *wdr41*^*−/−*^ brain and MEF cell lysates. (**G**) C9orf72 and SMCR8 levels were quantified and normalized to GAPDH for the experiment in (F) (n=3, *, p<0.05, student’s t-test).

### SMCR8 deficiency leads to reduced levels of proteins in the autophagy-lysosome pathway

Several studies have indicated a role for SMCR8 in the autophagy-lysosome pathway. C9orf72-SMCR8-WDR41 was reported to associate with the RB1CC1-ULK1 autophagy initiation complex [8, 10, 11]. More recently SMCR8 was shown to regulate *Ulk1* gene expression [11]. Thus we decided to determine autophagy-lysosome defects in fibroblasts derived from *smcr8*^*−/−*^ mice. To our surprise, we found that the levels of many proteins in the autophagy-lysosome pathway were significantly decreased in *smcr8*^*−/−*^ fibroblasts, including LC3B, SQSTM1, LAMP1, RB1CC1 and ULK1 (Fig. 3A-3D). Many of the corresponding genes are under the transcriptional control of TFEB, a master regulator of lysosomal biogenesis [23], indicating that TFEB might be affected by SMCR8 ablation. A significant reduction in TFEB protein levels was observed in *smcr8*^*−/−*^ fibroblasts (Fig. 3A), explaining the decreased levels of many autophagy-lysosomal proteins. Autophagy flux assay indicates SQSTM1 and LC3 protein levels were deceased in *smcr8*^*−/−*^ fibroblasts even with lysosomal activities inhibited (Fig. 4A-C) without changes in autophagy flux (Fig. 4D), indicating that the changes might occur at transcriptional levels. Indeed, qPCR analysis showed decreased mRNA levels of *Tfeb, Lamp, Sqstm1, Lc3b, Ulk1* and *Rb1cc1*in *smcr8*^*−/−*^ fibroblasts (Fig. 4E).

**Figure 3.**
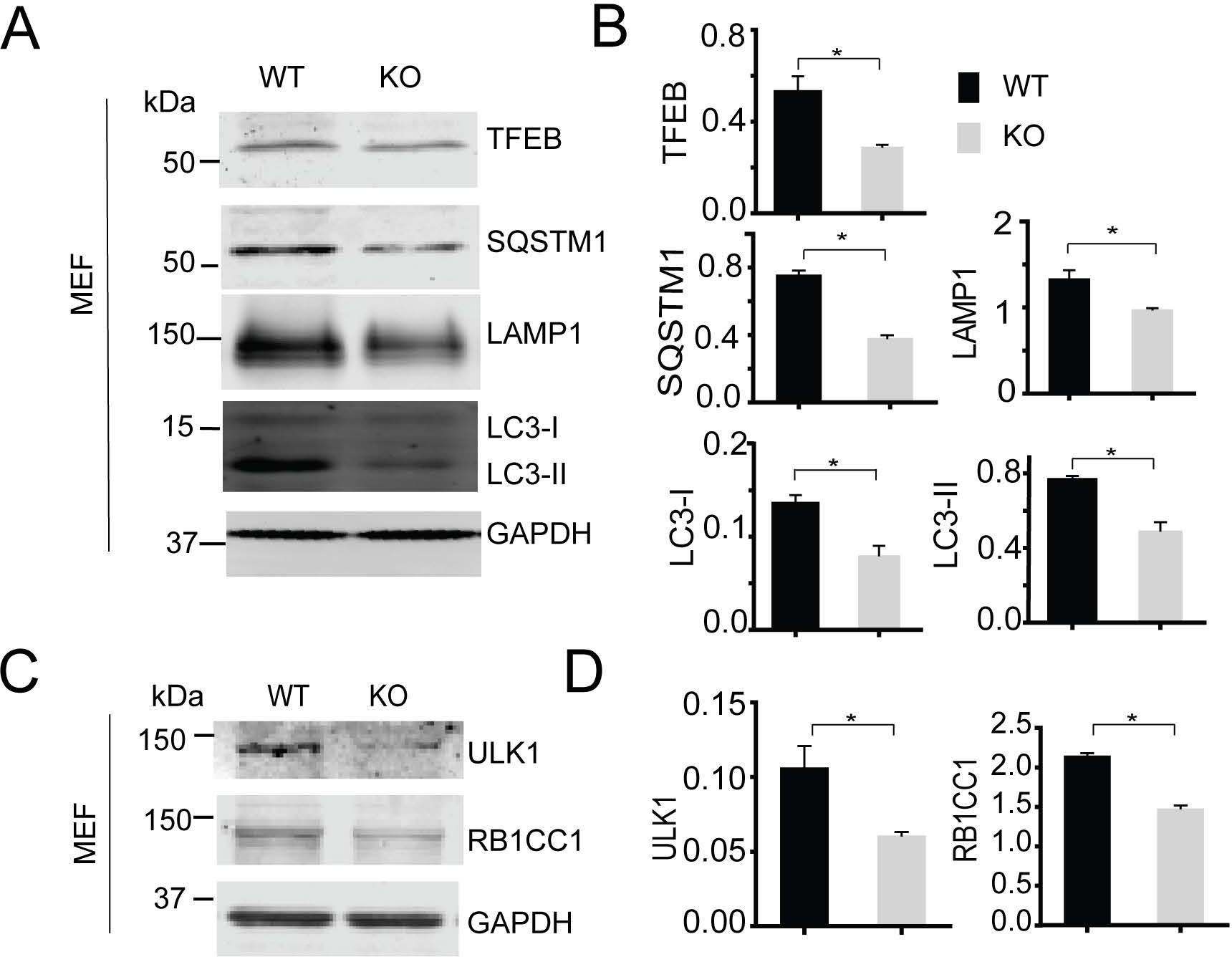
Reduced levels of TFEB and autophagy-lysosome proteins in *smcr8*^*−/−*^ fibroblasts. (**A, B**) Western blot analysis of TFEB, SQSTM1, LC3, LAMP1 and GAPDH proteins in WT and *smcr8*^*−/−*^ MEF lysates. Proteins levels were quantified and normalized to GAPDH (n=3, *, p<0.05, student’s t-test). (**C,D**) Western blot analysis of ULK1 and RB1CC1 proteins in WT and *smcr8*^*−/−*^ MEF lysates. Proteins levels were quantified and normalized to GAPDH (n=3, *, p<0.05, student’s t-test)

**Figure 4.**
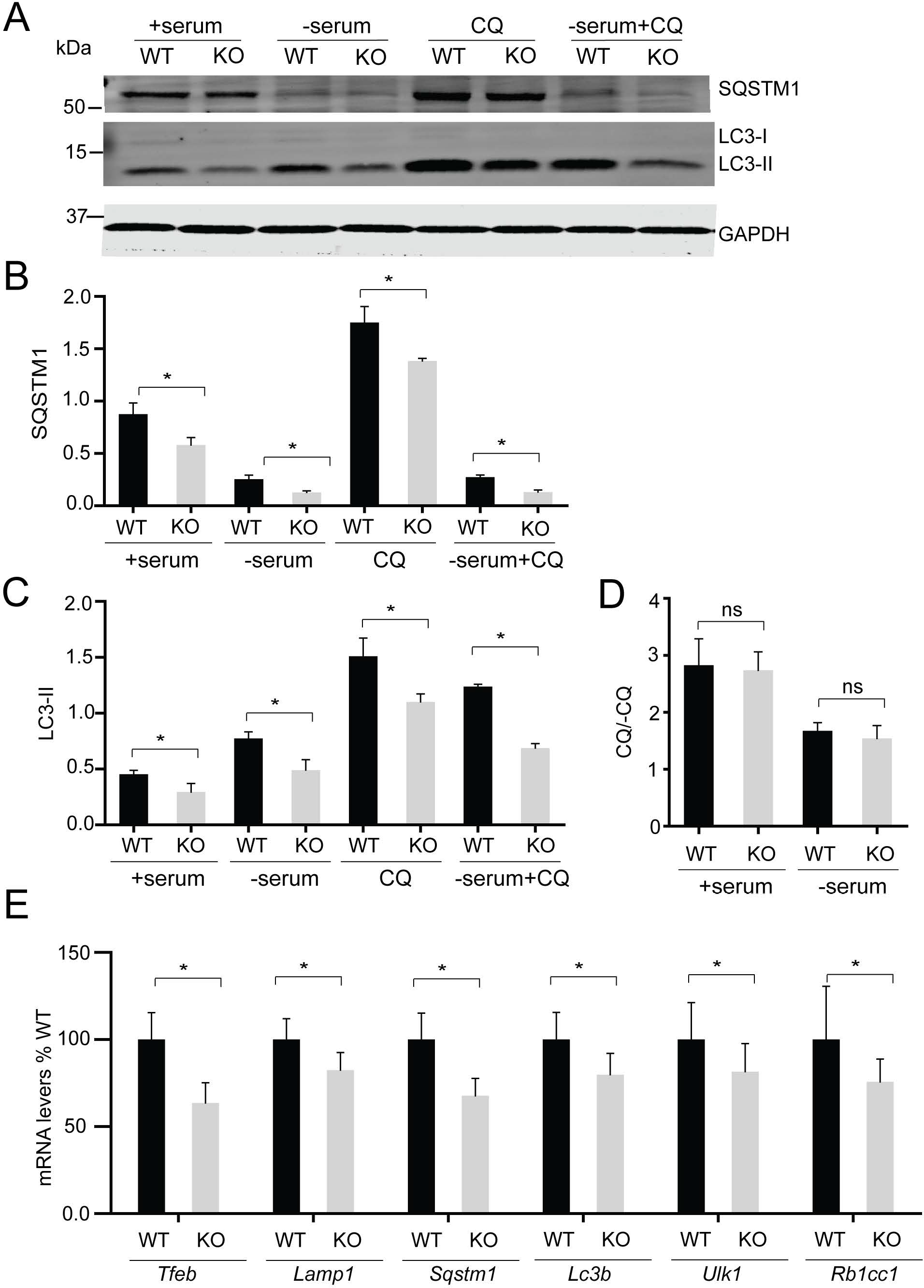
Analysis of autophagy flux and mRNA levels of autophagy-lysosome genes in *smcr8*^*−/−*^ fibroblasts. (**A-C**) Western blot analysis of SQSTM1, LC3, and GAPDH proteins in WT and *smcr8*^*−/−*^ MEF lysates under the indicated conditions. Cells were in 10% FBS or serum starved for 6 h with or without chloroquine treatment (CQ). Proteins levels were quantified and normalized to GAPDH (n=3, *, p<0.05, student’s t-test). (**D**) Autophagy flux was calculated by dividing the LC3-II:GAPDH in the presence of chloroquine by LC3-II:ACTB at baseline. (**E**) qPCR analysis of *Tfeb*, *Lamp1*, *Sqstm1*, *Lc3b*, *Ulk1* and *Rb1cc1* mRNA levels in WT and *smcr8*^*−/−*^ fibroblasts. The mRNA levels are normalized to *Actb* (n=3, *, p<0.05, student’s t-test).

Similar results were obtained from cortical neurons cultured from *smcr8*^*−/−*^ mouse pups. A significant decrease in the protein levels of TFEB accompanied by concomitant decreased levels of LAMP1, SQSTM1 and LC3B was detected in *smcr8*^*−/−*^ neurons compared to WT controls (Fig. 5A and B). The decrease in TFEB and SQSTM1 proteins was also observed in *smcr8*^*−/−*^ brain lysates compared to that of littermate WT controls (Fig. 5C and D).

**Figure 5.**
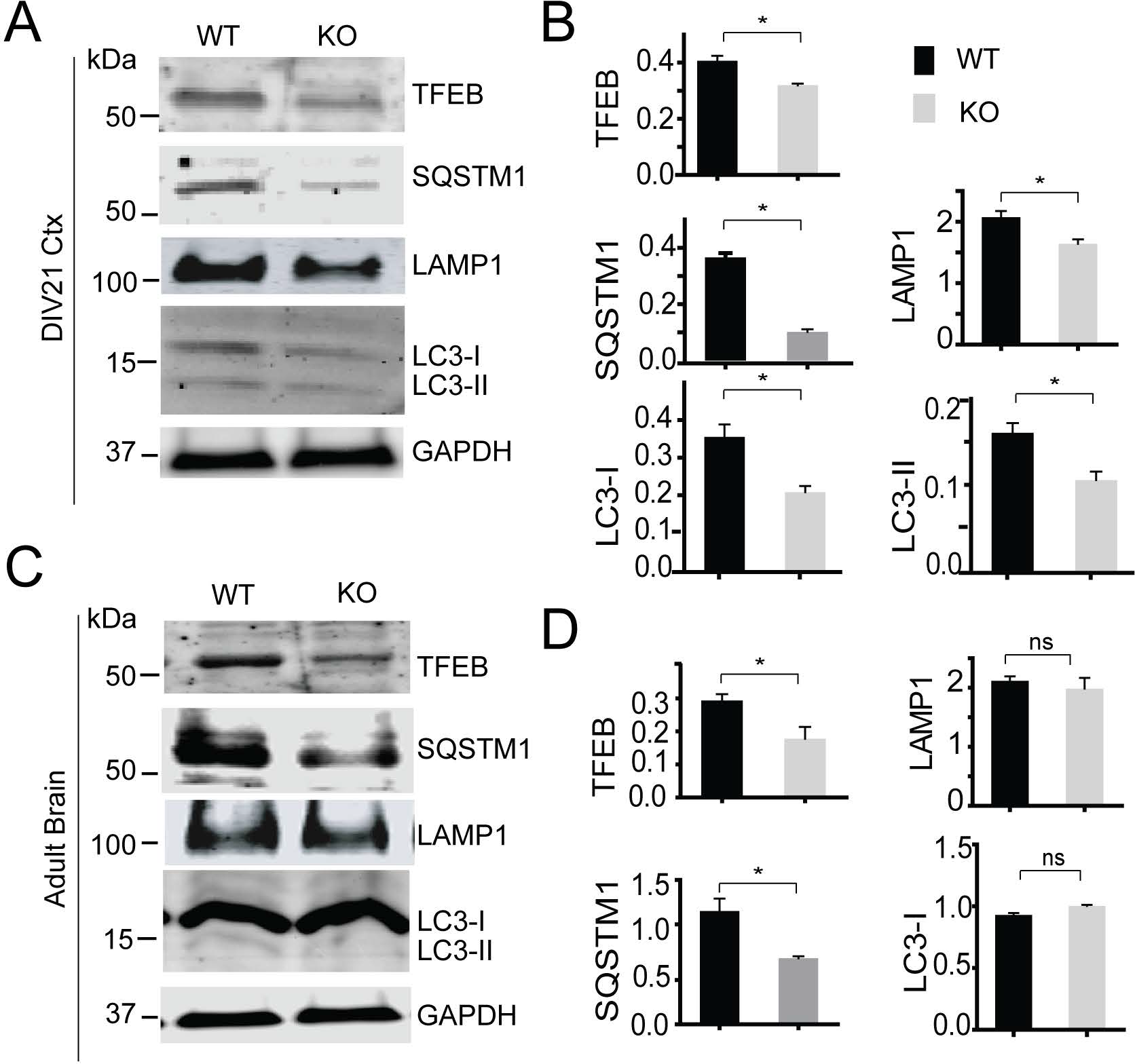
Reduced levels of TFEB and autophagy-lysosome proteins in *smcr8*^*−/−*^ neurons and brain lysates. (**A,B**) Western blot analysis of TFEB, SQSTM1, LC3, LAMP1 and GAPDH proteins in day in vitro (DIV)21 cortical neurons cultured from WT and *smcr8*^*−/−*^ pups. Proteins levels were quantified and normalized to GAPDH (n=3, *, p<0.05, student’s t-test). Ctx, cortical neurons. (**C,D**) Western blot analysis of TFEB, SQSTM1, LC3, LAMP1 and GAPDH proteins in brain lysates from 4-month-old WT and *smcr8*^*−/−*^ mice. Proteins levels were quantified and normalized to GAPDH (n=3; *, p<0.05; ns, not significant; student’s t-test).

Nuclear translocation of TFEB is required for its induction of lysosomal biogenesis and is regulated by nutrient deprivation [24, 25]. Immunostaining of TFEB revealed that not only the protein levels of TFEB were affected in *smcr8*^*−/−*^ cells, but also TFEB subcellular localization. In *smcr8*^*−/−*^ fibroblasts, TFEB remained cytoplasmic even upon serum starvation(Fig. 6A and B) and less nuclear TFEB was observed in *smcr8*^*−/−*^ primary neurons compared to that of WT controls (Fig. 6C, 6D). Under conditions of amino acid starvation, TFEB was translocated to the nucleus in *smcr8*^*−/−*^ fibroblasts, yet there was a small but significant decrease in TFEB nuclear signal (Fig. 6A and B), which might be due to reduced TFEB protein levels in *smcr8*^*−/−*^ fibroblasts.

**Figure 6.**
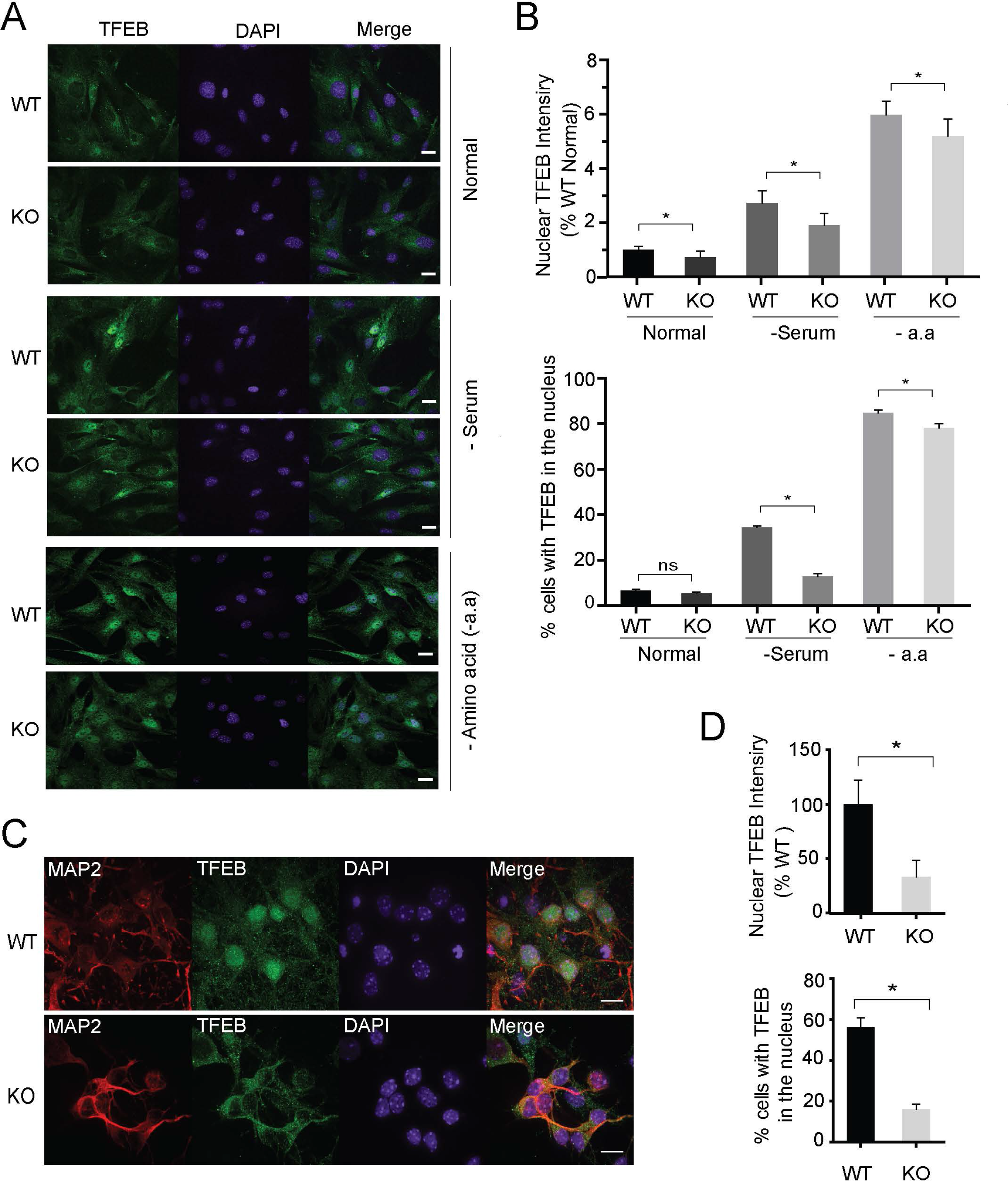
TFEB nuclear translocation is impaired in *smcr8*^*−/−*^ MEFs and neurons. (**A, B**) WT and *smcr8*^*−/−*^ MEFs were either grown in serum-containing medium (+serum), or starved for 4 h in DMEM (-serum) or 2 h in RPMI medium without amino acids (-a.a). Cells were fixed and stained with rabbit anti-TFEB antibodies and Hoechst. Scale bar: 10 μm. Nuclear and cytoplasmic TFEB signals were analyzed using Image J software. At least 50 cells were analyzed in each treatment. (**C, D**) WT and *smcr8*^*−/−*^ DIV7 cortical neurons were fixed and stained with rabbit anti-TFEB antibodies, mouse anti-MAP2 antibodies and Hoechst. Scale bar: 10 μm. Nuclear and cytoplasmic TFEB signals were analyzed using ImageJ software.

### Increased MTORC1 and AKT activity due to SMCR8 deficiency

It is well established that TFEB nuclear translocation is regulated by the MTORC1 kinase [24, 25]. In growth conditions, TFEB is phosphorylated by MTORC1 and phosphorylated TFEB is recognized by the YWHA/14-3-3 protein to sequester TFEB in the cytosol [24, 25]. Thus, reduced TFEB nuclear localization in *smcr8*^*−/−*^ cells prompted us to determine if there were any changes in MTORC1 activity. Indeed MTORC1-dependent phosphorylation of RPS6KB1/S6K (ribosomal protein S6 kinase, polypeptide 1) was significantly increased in *smcr8*^*−/−*^ fibroblasts and neurons compared to WT controls (Fig. 7A and B), indicating elevated MTORC1 activation in *smcr8*^*−/−*^ cells. MTORC1 activation requires recruitment to lysosomal membrane as well as phosphorylation by the upstream kinase AKT [24, 25]. A significantly increased colocalization between MTORC1 and the lysosomal membrane protein LAMP1 was detected in *smcr8*^*−/−*^ fibroblasts and neurons (Fig. 7C-7F). Activation of RRAGs/Rag GTPases by amino acids is essential for lysosomal recruitment of MTOR [24, 25]. We found that amino acid withdrawal led to dissociation of MTORC1 from lysosomes in *smcr8*^*−/−*^ cells (Fig. 7C and D), indicating that lysosomal recruitment and activation of MTORC1 in *smcr8*^*−/−*^ cells remains amino-acid dependent.

**Figure 7.**
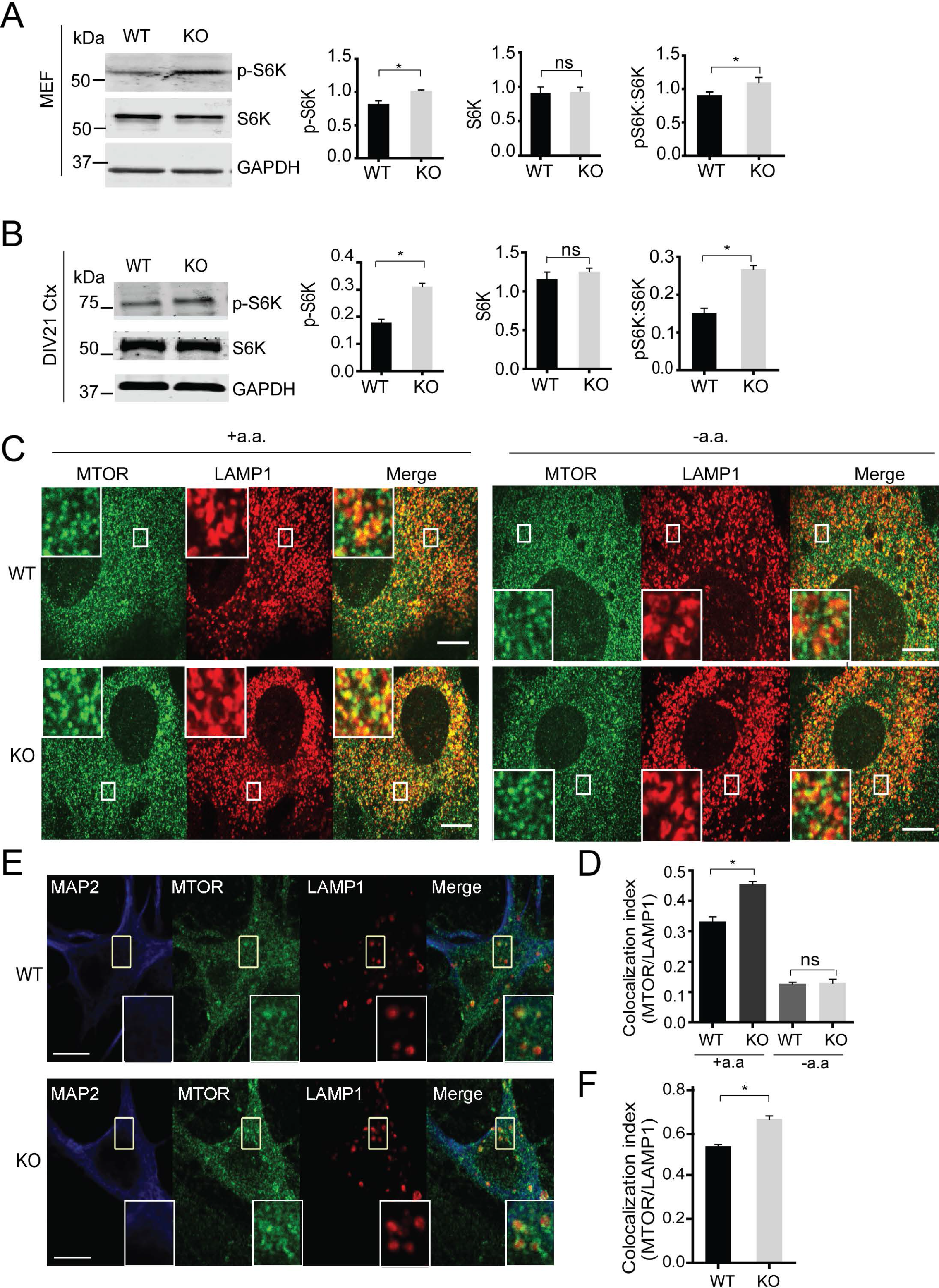
Increased MTORC1 activities in *smcr8*^*−/−*^ cells. (**A,B**) Western blot analysis of p-RPS6KB/S6K, RPS6KB/S6K and GAPDH in WT and *smcr8*^*−/−*^ fibroblasts lysates (**A**) or DIV21 cortical neuron lysates (**B**). p-RPS6KB/S6K and RPS6KB/S6K proteins levels were quantified and normalized to GAPDH (n=3, *, p<0.05, student’ s t-test). Ctx, cortical neurons. (**C, D**) Immunostaining of MTORC1 and LAMP1 and quantification of MTORC1 and LAMP1 colocalization (Manders’ Coefficient) using ImageJ in WT and *smcr8*^*−/−*^ MEFs with or without amino acid starvation (n=3; *, p<0.05; ns, not significant; student’s t-test). (**E, F**) Immunostaining of MTORC1 and LAMP1 and quantification of MTORC1 and LAMP1 colocalization using ImageJ in DIV21 WT and *smcr8*^*−/−*^ cortical neurons (n=3; *, p<0.05; ns, not significant; student’s t-test). Scale bar: 10 μm.

To determine whether AKT activities are altered in *smcr8*^*−/−*^ cells, we first examined the ratio of phosphorylated MTOR (S2448) to total MTOR and found significantly increased MTOR phosphorylation in *smcr8*^*−/−*^ cells (Fig. 8A-8D). Furthermore, the phosphorylation of AKT at Ser473 and Thr308, mediated by MTORC2 and PDK1 upon growth factor stimulation, was significantly increased in *smcr8*^*−/−*^ fibroblasts, cortical neurons, and brain lysates compared to those of WT controls (Fig. 8E-8J). Importantly, increased AKT activation in *smcr8*^*−/−*^ fibroblasts could be rescued by viral expression of GFP-tagged SMCR8 (Fig. 9A and B). These data support the idea that SMCR8 acts as a negative regulator of AKT-MTORC1 signaling. Treatment with the phosphatidylinositol 3-kinase (PtdIns3K) inhibitor LY294002 abolished increased AKT phosphorylation in *smcr8*^*−/−*^ fibroblasts (Fig. 9C and D), indicating that elevated PtdIns3K activities drive AKT-MTORC1 signaling in *smcr8*^*−/−*^ cells.

**Figure 8.**
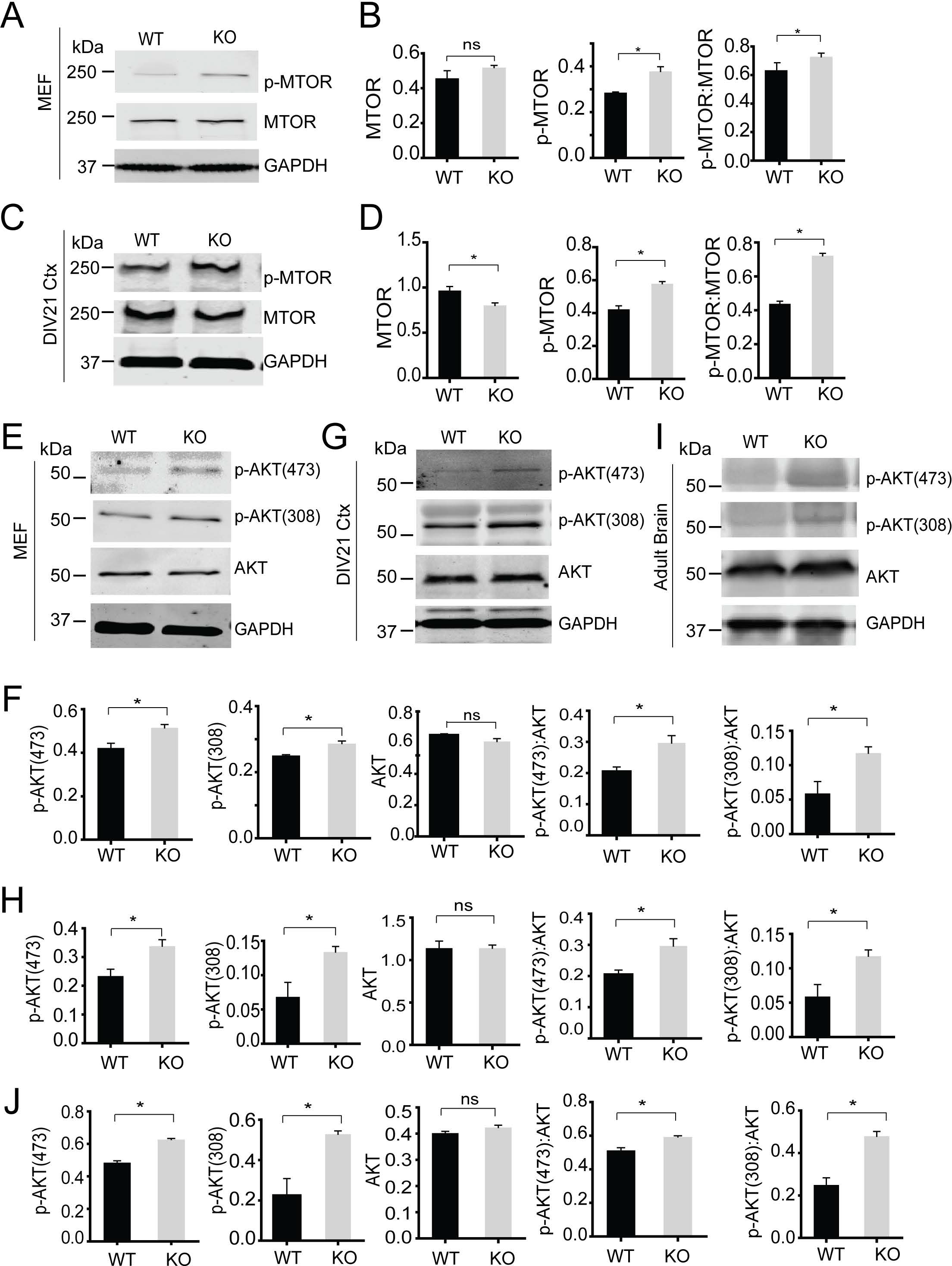
Increased AKT signaling due to SMCR8 deficiency. (**A-D**) Western blot analysis of p-MTOR, MTOR and GAPDH in WT and *smcr8*^*−/−*^ MEF lysates (**A,B**) and DIV21 cortical neurons (**C,D**). p-MTOR and MTOR proteins levels were quantified and normalized to GAPDH (n=3; *, p<0.05; ns, not significant; student’s t-test). (**E-J**) Western blot analysis of p-AKT T308, p-AKT S475, AKT and GAPDH in WT and *smcr8*^*−/−*^ MEF lysates (E,F), DIV21 cortical neurons (G,H) and adult brain lysates (I,J). p-AKT and AKT proteins levels were quantified and normalized to GAPDH (n=3; *, p<0.05; ns, not significant; student’s t-test).

**Figure 9.**
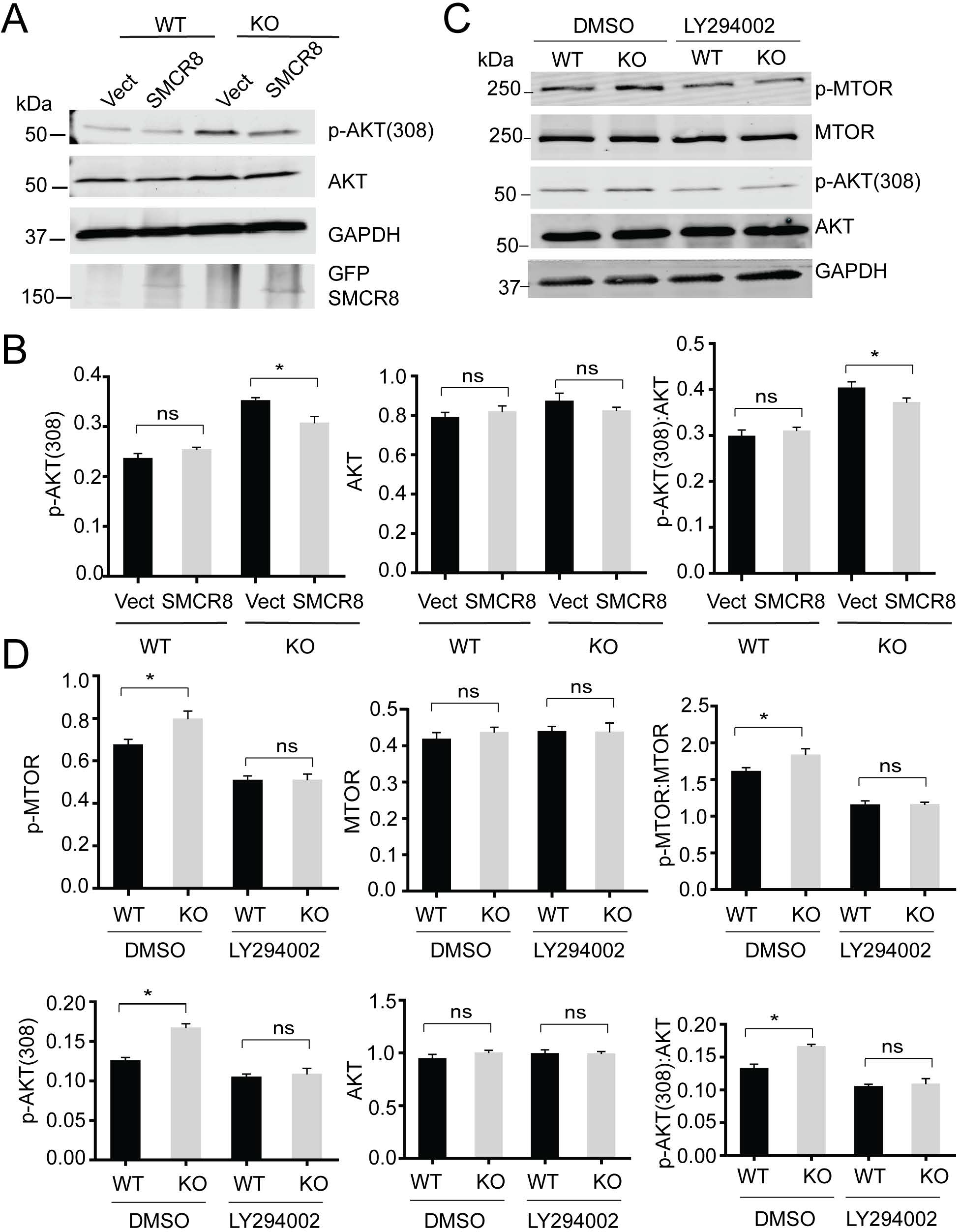
PtdIns3K inhibition rescues increased AKT activities in SMCR8-deficient fibroblasts. (**A,B**) WT and *smcr8*^*−/−*^(KO) fibroblasts were infected with control lentiviruses or lentiviruses expressing GFP-SMCR8. The levels of AKT and p-AKT (308) were analyzed and normalized to GAPDH (n=3; *, p<0.05; ns, not significant; student’s t-test). (**C, D**) WT and *smcr8*^*−/−*^ (KO) fibroblasts were treated with DMSO or PtdIns3K inhibitor LY294002 (25 μM) for 4 h. The levels of p-MTOR, MTOR, AKT and p-AKT (308) were analyzed and normalized to GAPDH (n=3, *, p<0.05, student’s t-test).

### SMCR8 deficiency leads to defects in synaptic pruning

MTORC1 and autophagy activities are tightly linked to synaptic plasticity and spine morphogenesis [26]. Because we have observed increased MTORC1 and AKT activities and lower levels of autophagy-lysosomal proteins in *smcr8*^*−/−*^ cortical neuron and brain lysates, we examined changes in spine density upon SMCR8 loss. Western blot analysis of DLG4/PSD95, a postsynaptic marker, revealed a significant increase in DLG4 protein levels in lysates derived from *smcr8*^*−/−*^ brain or cortical neuron cultures compared to those from littermate WT controls, while the levels of the presynaptic marker, SYN1 (synapsin I), were not changed (Fig. 10A and B). Consistent with this, Golgi staining of brain sections revealed a significant increase in spine density in the primary auditory cortex in *smcr8*^*−/−*^ mice (Fig. 10C). This result could be caused by increased spine growth or reduced spine pruning due to the increased MTORC1 activities in *smcr8*^*−/−*^ mice.

**Figure 10.**
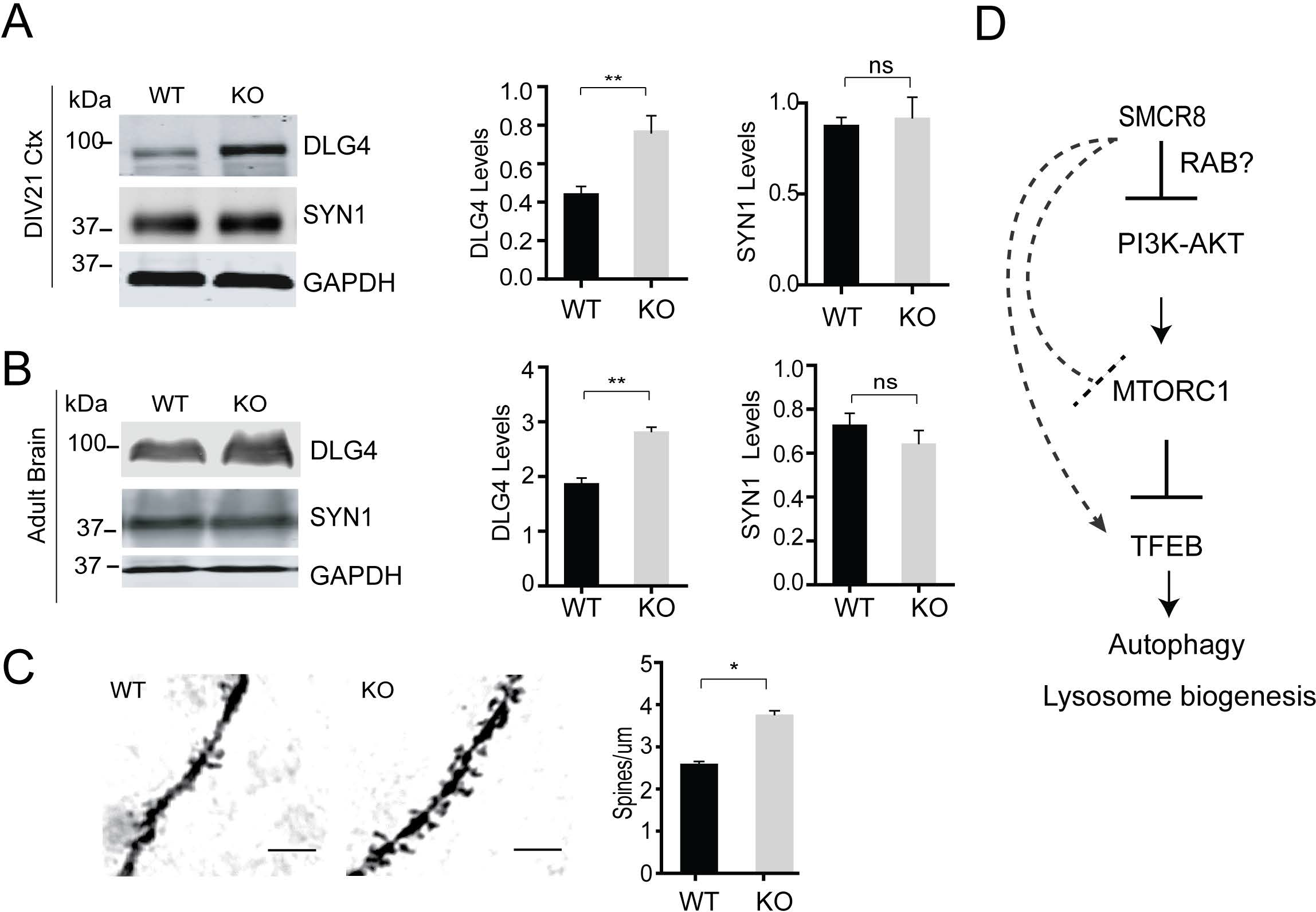
SMCR8 deficiency results in increased spine density. (**A,B**) Western blot analysis of PSD95, SYN1 and GAPDH in WT and *smcr8*^*−/−*^ DIV21 cortical neurons (**A**) and brain lysates (**B**). DLG4/PSD95 and SYN1 (synapsin I) proteins levels were quantified and normalized to GAPDH (n=3; *, p<0.05; ns, not significant; student’s t-test). Ctx, cortical neurons. (**C**) Golgi staining of primary auditory cortexes in 6-months-old WT and *smcr8*^*−/−*^ mice. The number of spines along dendrites was quantified. Scale bar: 10 μm. n=3, *, p<0.05, student’s t-test. (**D**) SMCR8 negatively regulates AKT and MTORC1 signaling to modulate autophagy and lysosome activities. Increased AKT and MTORC1 activities due to SMCR8 deficiency leads to decreased autophagy activities and lysosomal biogenesis and subsequent tissue homeostasis defects. SMCR8 might also regulate lysosomal recruitment of MTOR and *Tfeb* transcription through other unknown mechanisms.

## Discussion

### SMCR8 negatively regulates MTORC1 and AKT signaling

Our data supported the hypothesis that SMCR8 negatively regulates AKT and MTORC1 signaling to modulate lysosome biogenesis and tissue homeostasis (Fig. 10D). Increased AKT activity due to SMCR8 deficiency results in enhanced MTORC1 signaling and significant reduction in the protein levels of autophagy-lysosome proteins in cortical neurons and fibroblasts. However, the mechanism by which SMCR8 regulates AKT remains to be elucidated. Since SMCR8 contains DENN domains [13, 14], we suspect that SMCR8 might regulate AKT activities through RAB GTPases. Recently, SMCR8 and C9orf72 have been shown to function as a GEF for RAB5, RAB7, RAB11, RAB8A and RAB39B [7, 10, 15, 16], and might also function as the effector for RAB1 [27]. Since there are more than 70 RAB GTPases in the human genome, it is highly possible that there are additional RABs regulated by SMCR8 that affect AKT signaling.

It is intriguing that both C9orf72 and SMCR8 contain a DENN domain, and these 2 proteins form a tight complex. This mimics the interaction between FLCN (folliculin) and FNIP1 (folliculin interacting protein 1), and between NPR2 (natriuretic peptide receptor 2), and NPR3. Interestingly, the FLCN-FNIP1 complex was shown to function as the GAP for the RRAGC/D GTPases involved in recruitment of MTORC1 to lysosomal membrane [28]. Thus, by analogy, there is a possibility that the C9orf72-SMCR8 complex might also function as a GAP instead of a GEF for RAB GTPases. Future in vitro GEF and GAP activity assays with purified C9orf72 and SMCR8 recombinant proteins and RAB GTPases and determination of RAB activity changes in cells with C9orf72 or SMCR8 deficiency will allow us to define the role of C9orf72 and SMCR8 in RAB GTPase regulation.

Although AKT and MTORC1 activities are significantly increased when SMCR8 is lost, we failed to detect an increase in AKT and MTORC1 activities in C9orf72-deficient fibroblasts (data not shown), indicating that SMCR8 is likely to play a more critical role in regulating AKT-MTORC1 signaling than C9orf72. However, this is very puzzling because SMCR8 protein levels were drastically reduced in *c9orf72* knockout (Fig. 2). It is possible that the residual SMCR8 proteins in C9orf72-deficient cells are sufficient to restrict AKT regulation or, alternatively, in the absence of SMCR8, the function of C9orf72 is altered. Fibroblasts deficient in both C9orf72 and SMCR8 show enhanced AKT activities similar to that of SMCR8 ablation (data not shown), arguing against the latter possibility.

Increased localization of MTOR to lysosomal membrane indicates SMCR8 might also affect lysosomal recruitment of MTOR, which is mainly regulated by the amino acid sensing pathways mediated by the RRAG GTPases [28]. However, neither C9orf72 nor SMCR8 interacts with RRAG GTPases (data not shown). In SMCR8-deficient fibroblasts, MTOR still dissociates from the lysosome membrane upon amino acid starvation, suggesting that SMCR8 might regulate MTOR lysosomal recruitment but it is not an essential inhibitor of the pathway (Fig. 10D).

### SMCR8 and lysosome biogenesis

The levels of many autophagy-lysosomal proteins are significantly decreased in SMCR8-deficient cells. qPCR analysis revealed that this likely happened at transcriptional levels (Fig. 4D). Intriguingly, both mRNA and protein levels of TFEB were decreased in *smcr8*^*−/−*^ cells (Fig. 3A and Fig. 4D). This could be due to decreased TFEB nuclear translocation and reduced self-induction of its mRNA expression by TFEB in *smcr8*^*−/−*^ cells, or, alternatively, SMCR8 might regulate TFEB transcription and protein levels through another unknown mechanism (Fig. 10D).

### SMCR8 regulates spine density

Another interesting phenotype of SMCR8-deficient mice is increased spine density compared to that of littermate WT controls. MTORC1 regulates spine growth through protein synthesis and regulates spine pruning through autophagy [26]. Thus, it is not surprising that increased MTORC1 activities in the *smcr8*^*−/−*^ mice will lead to greater spine density. However, it is still not clear whether this is due to increased growth or diminished spine pruning. Also, it is unclear whether microglia plays a role in this regulation. It will be interesting to test whether C9orf72 haploinsufficiency due to hexanucleotide expansion leads to changes in SMCR8 protein levels and subsequently spine density and whether that plays a role in ALS-FTLD disease progression.

### C9orf72-SMCR8 and immune regulation

The *in vivo* phenotypes of SMCR8 deficiency are highly similar to that of C9orf72 deficiency with splenomegaly, inflammation in multiple tissues and autoimmunity, suggesting that C9orf72 and SMCR8 function together as a complex in the hematopoietic system to restrict autoimmunity [29]. MTORC1 is a critical regulator of immune responses, and increased MTORC1 activities trigger autoimmunity [30]; thus, it is highly possible that C9orf72-SMCR8 acts through MTORC1 to regulate immune responses. Although *C9orf72* mRNA and protein levels are decreased in ALS-FTLD patients with hexanucleotide expansions, the prevailing disease mechanisms are gain of toxicity from the repeat expansions [31]. However, autoimmunity strongly associates with ALS-FTLD [32]. Although microglia do not show any obvious morphology changes in the *c9orf72* knockout or *Smcr8* knockout mice under physiological conditions, it remains to be tested whether C9orf72 and SMCR8 regulate microglial function under challenge, and whether haploinsufficiency of C9orf72 could affect disease progression through regulating microglial behavior and brain inflammation.

## Materials and Methods

### Ethics statement

The animal protocol (2017-0056) was approved by the Cornell University animal care and use committee following the National Research Council’s guide to the care of laboratory animals.

### Pharmacological reagents and antibodies

The following primary antibodies were used in this study: Anti-phospho-T308 AKT (sc-271966) and phospho-S475 AKT (sc-81433), anti-AKT (sc-81434), and anti-ULK1 (sc-33182) from Santa Cruz Biotechnology, anti-C9orf72 (AP12928b), mouse anti-GAPDH (60004-1-1g), anti-RPS6KB/S6K (14485-1-AP), anti-SYN1 (17785-1-AP) and anti-RB1CC1 (17250-1-AP) from Proteintech, anti-LC3 (PM036) from MBL international, anti-SQSTM1 (NBP1-42821) from Novus, anti-MTORC1 (7C10), anti-phospho-MTORC1 (Ser2448), anti-p-RPS6KB/S6K (9234S) from Cell Signaling Technology, rabbit anti-IBA1 from Wako, mouse anti-MAP2 (556320) and rat anti-mouse LAMP1 (553792) from BD Biosciences, sheep anti-GRN/progranulin antibodies from R&D systems (AF2557), anti-DLG4/PSD95 (2868029) from EMD Millipore Crop and anti-SMCR8 antibodies from Thermo Fisher Scientific (TA2502611D). The following secondary antibodies were used: donkey anti-mouse Alexa Fluor 800 (926-32212) and donkey anti-rabbit Alexa Fluor 800 (926-32213) from LI-COR, donkey anti-goat Alexa Fluor 680 (A32860), donkey anti-rabbit Alexa Fluor 680 (A10043), donkey anti-mouse Alexa Fluor 680 (A10038), and donkey anti-rat Alexa Fluor 680 (A21096), donkey anti-mouse Alexa Fluor 594 (A21203), donkey anti-rabbit Alexa Fluor 488 (A21206), donkey anti-mouse Alexa Fluor 647 (A31571), donkey anti-sheep Alexa Fluor 647 (A21448) and Hoechst stain (H1339) from Thermo Fisher Scientific. Chloroquine (C-6628) is from Sigma-Aldrich and PtdIns3K inhibitor LY294002 (L-7962) is from LC laboratories.

### Mouse strains

*smcr8* knockout mice were produced using CRISPR-Cas9 genome editing with a guide RNA (gRNA, sequence: GATCAGCGCCCCTGATGTGG) targeting near the start codon of the mouse *Smcr8* gene. C57BL/6J × FvB/N mouse embryos were injected with gRNA and *Cas9* mRNA at the Cornell Transgenic Core Facility. Editing was confirmed by sequencing PCR products from genomic DNA. Offspring from the founder containing a 128-base pair (bp) deletion were used for the study. The following primers were used to genotype *smcr8* knockout mice: 5’-GCTGGTGACCTAGCTTCAGG-3’ (forward) and 5’-ACCGACATAATCCGCAAAGA-3 (reverse) (PCR product: 595 bp). *wdr41* knockout mice were created in a similar manner with the guide RNA (5’-AGCCGAGGTAATGCGCGGGGG-3’) resulting in a 122-nucleotide deletion shortly after the start-codon.

### Cell culture

Primary cortical neurons were isolated from P0-P1 pups using a modified protocol [33]. Cortices were rapidly dissected from the brain in 2 mL HBSS (Corning Cellgro, 23-21-020-CV) supplemented with B27 (Thermo Fisher Scientific, 17504044) and 0.5 mM L-glutamine (Thermo Fisher Scientific, 25030-081) at 4°C. Meninges and excess white matter were removed before digestion with papain (Worthington, LS003119; 2 mg/ml in HBSS) and DNaseI (1 mg/ml in HBSS; Sigma-Aldrich, D4513) for 12 min at 37°C. Tissues were then dissociated using fire-polished glass pipettes. Cells were spun down and resuspended in Neurobasal-A medium (Thermo Fisher Scientific, 10888022) plus B27 (Thermo Fisher Scientific, 17504044) and plated onto poly-lysine (Sigma-Aldrich, 27964-99-4)-coated dishes.

Primary mouse fibroblasts were isolated from mouse newborn pups using collagenase type II (Worthington, LS004204) and grown in DMEM plus 10% fetal bovine serum. For all the western blot and qPCR studies, passage 7 to 10 primary fibroblasts were used.

### Protein analysis

Cells and tissues were lysed in RIPA buffer (50 mM Tris, pH 8.0, 150 mM NaCl, 1% Triton X-100(Sigma-Aldrich, 78787), 0.1% SDS, 0.1% deoxycholic acid (Sigma-Aldrich, 83-44-3) with protease and phosphatase inhibitors (Sigma-Aldrich, P5726). Samples were denatured in 2xSDS sample buffer (4% SDS, 20% glycerol, 100 mM Tris, pH 6.8, 0.2 g/L bromophenol blue) by boiling for 3 min. Samples were run on 8%, 12% or 15% polyacrylamide gels and transferred to PVDF membranes (Millipore, IPFL00010). Membranes were blocked in either Odyssey Blocking Buffer (LI-COR Biosciences, 927-4000) or 5% non-fat milk in PBS (Gibico, 21600-044) for 1 h followed by incubation with primary antibodies overnight at 4°C. Membranes were washed 3 times with Tris-buffered saline (Amresco,0497) with 0.1% Tween-20 (Amresco, 0777) (TBST) then incubated with secondary antibodies for 2 ho at room temperature. Membranes were washed 3 times with TBST and imaged using an Odyssey Infrared Imaging System (LI-COR Biosciences) with settings in the linear range of band intensities. Western blot signals were then analyzed using the ImageJ software.

### Hematoxylin and eosin (H&E) staining

Mouse tissues were fixed with 4% paraformaldehyde (Sigma-Aldrich, P6148). After dehydration with 70% ethanol, tissues were embedded with paraffin. The tissues were sliced to 8 μm. Following deparaffiinization with xylene and ethanol (100%, 95%, 80%) and rehydration with tap water, the slides were stained in hematoxylin for 3 min, destained with acid ethanol and rinsed with tap water, and then stained with eosin for 30 s. The slides were then dehydrated with ethanol and xylene and mounted.

### Golgi staining

For Golgi-Cox staining, 6-month-old mice were sacrificed by rapid decapitation and their brains were removed. The brains were immersed in FD Rapid Golgi Stain™ kit solution (FD Neurotechnologies, PK401) and processed according to the manufacturer’s instructions. Then, the tissues were sliced to 60 μmand stained according to the manufacturer’s protocol. For quantification of dendritic spines, primary auditory cortexes were imaged using a CSU-X spinning disc confocal microscope (Intelligent Imaging Innovations) with an HQ2 CCD camera (Photometrics) using a 100x objective. Four or 5 dendritic segments per cell were analyzed and the spine density is presented as the mean number of spines per 10 μm of dendritic length, calculated using a double-blind method. Experiments were independently repeated with 3 mouse brains each group.

### ELISA

Serum samples were collected from adult mice and analyzed using a mouse anti-dsDNA Ig’s (Total A+G+M) ELISA Kit from Alpha Diagnostic Interanational (5110) according to the manufacturer’s instructions.

### Immunofluorescence microscopy

Cells grown on glass coverslips were fixed in 3.7% paraformaldehyde for 15 min, washed 3 times with PBS, and permeabilized and blocked in Odyssey Blocking Buffer with 0.05% saponin (Alfa Aesar, AI8820) or 0.1% Triton X-100 (Sigma-Aldrich, 78787) for 20 min. Primary antibodies diluted in blocking buffer with 0.05% saponin were applied to the cells overnight at 4°C. Coverslips were washed 3 times with PBS. Secondary antibodies and Hoechst stain diluted in blocking buffer with 0.05% saponin were applied to the cells for 2 h at room temperature. Coverslips were washed and mounted onto slides with Fluoromount G (Southern Biotech, 0100-01). Images were acquired on a CSU-X spinning disc confocal microscope (Intelligent Imaging Innovations) with an HQ2 CCD camera (Photometrics) using a 100x objective. Mouse tissues were perfused and fixed with 4% formaldehyde. After gradient dehydration with 15% and 30% sucrose (Amresco, 57-50-1), tissues were embedded with OCT compound (Sakura Finetek USA, 4583) and sectioned with a cryotome. For the immunostaining, tissue sections were permeabilized and blocked in Odyssey blocking buffer with 0.05% saponin for 1 h. Primary antibodies were incubated in blocking buffer overnight at 4°C. Sections were washed and incubated in secondary antibodies conjugated to Alexa Flour 488, 568, or 660 (Thermo Fisher Scientific, A11015, A11004, A21074). Sections were washed 3 more times and coverslips mounted onto slides with Fluoromount G (Southern Biotech, 0100-01). Images were acquired on a CSU-X spinning disc confocal microscope (Intelligent Imaging Innovations) with an HQ2 CCD camera (Photometrics) using a 40x objective.

### Image analysis

ImageJ software (NIH) was used to process and analyze the images. For TFEB nuclear translocation, nuclei were segmented using the blue channel and the cytoplasm was segmented using the green channel. The script calculates the ratio value resulting from the average intensity of nuclear green fluorescence divided by the average of the cytosolic intensity of green fluorescence. For each image, At least 50 cells were analyzed in each treatment. For MTOR and LAMP1 colocalization, background was subtracted and the colocalization between the LAMP1 and MTOR fluorescence was determined by Manders’ Coefficients using Plugins ‘JACOP’ of ImageJ software.

### RT-PCR analysis

RNA was purified from brain or fibroblasts using TRIzol Reagent (Thermo Fisher Scientific, 15596026). Two micrograms of total RNA were reverse transcribed using a poly(T) primer and SuperScript III Reverse Transcriptase (Thermo Fisher Scientific, 4368814). qPCR was performed on a LightCycler 480 (Roche Applied Science), and transcript levels were calculated using efficiency-adjusted ΔΔ-CT. All transcripts were normalized to *Actb*. The primers used are listed in Table S1.

### Statistical analysis

The data were presented as mean ± SEM. Two-group analysis was performed using the Student’s t test. P-values <0.05 were considered statistically significant.

## Supporting information

## Acknowledgements

We thank Ms. Xiaochun Wu for technical assistance, Teresa L. Southard for help with pathology and Dan Xia for help with ELISA and tissue processing. This work is supported by funding to F.H. from the Weill Institute for Cell and Molecular Biology, NINDS (R01NS088448, R01NS095954) and by funding to P.M.S. from the Harry and Samuel Mann Outstanding Graduate Student Award.

## List of abbreviations

ALS: amyotrophic lateral sclerosis
C9orf72: chromosome 9 open reading frame 72
FTLD: frontotemporal lobar degeneration
GEF: guanosine nucleotide exchange factor
GTPase: guanosine tri-phosphatase
KO: knockout
MTOR: mechanistic target of rapamycin kinase
SMCR8: Smith-Magenis chromosome region, candidate 8
WDR41: WD repeat domain 41
WT: wild type

