## Supplementary material for "SMCR8 negatively regulates AKT and MTORC1 signaling to modulate lysosome biogenesis and tissue homeostasis"

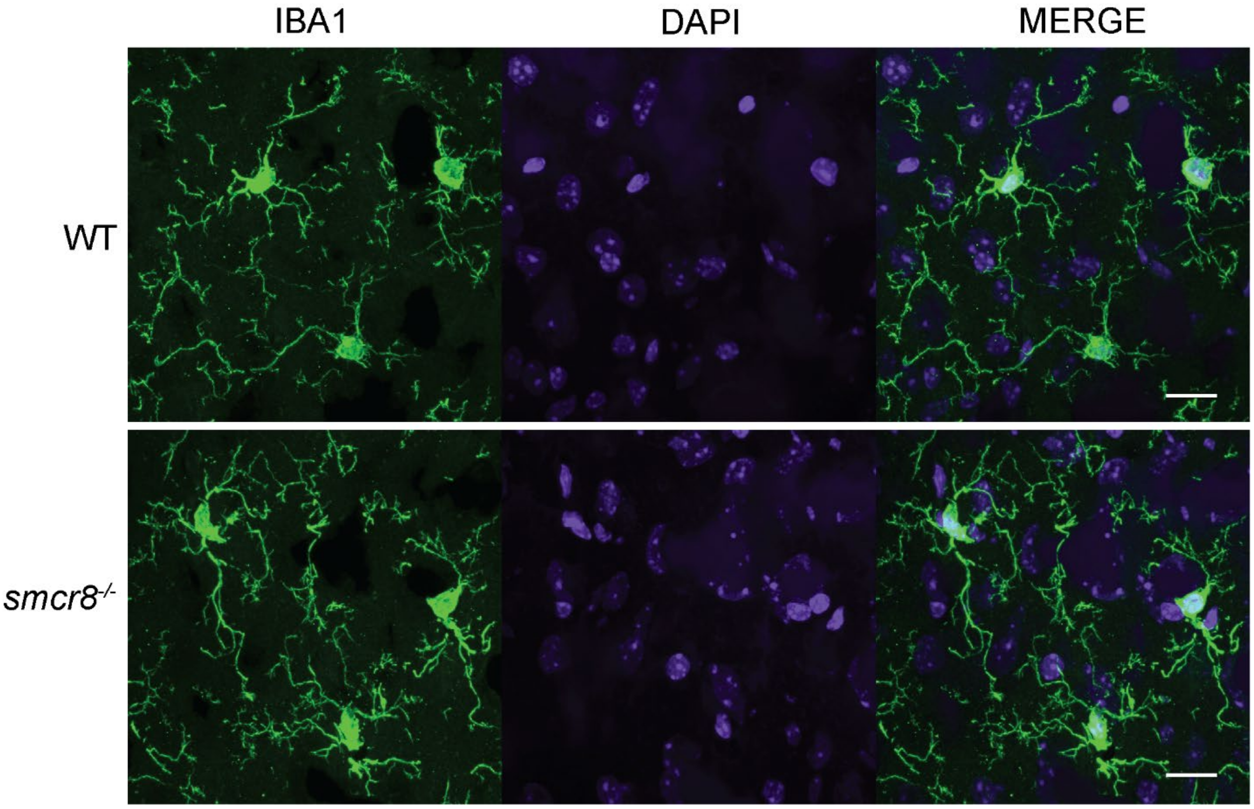


**Figure S1**. Microglia do not show any obvious phenotypes in SMCR8 deficient mice. Brain sections from 4 months old WT and *smcr8*$^{-/-}$ mice were stained with rabbit anti-IBA1 antibodies and Hoechst (to label nuclei). Scale bar:10µm.

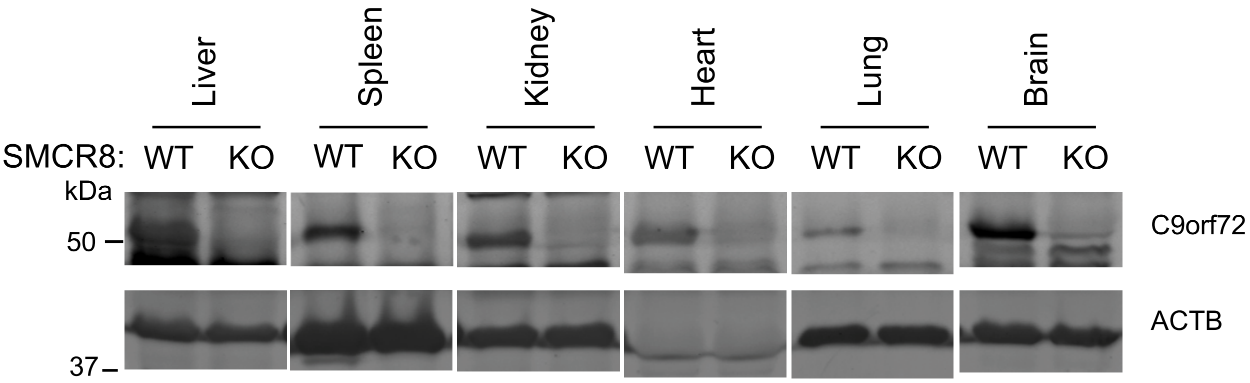


**Figure S2**. Decreased levels of C9orf72 protein in *smcr8*$^{-/-}$ tissue lysates. C9orf72 protein was analyzed in WT and *smcr8*$^{-/-}$ (KO) tissue lysates as indicated. ACTB was used as a loading control.

**Table S1**. Primers used for qPCR analysis in this study.

| Name | | Primer |
|---|---|---|
| *Actb* | forward | 5'- ACGAGGCCCAGAGCAAGAG -3' |
| | reverse | 5'- TCTCCAAGTCGTCCCAGTTG -3' |
| *C9orf72* | forward | 5'- GTCTTGGCAACAGCAGGAGAT -3' |
| | reverse | 5'- AGCAATCTCTGTCTTGGCAAC -3' |
| *Lamp1* | forward | 5'-TAATGGCCAGCTTCTCTGCCTCCT -3' |
| | reverse | 5'-AGGCTGGGGTCAGAAACATTTTCTT-3' |
| *Tfeb* | forward | 5'-GAGCTAACAGATGCTGAGAGCAGAGC-3' |
| | reverse | 5'-GCATCCTCCGGATGTAATCCACAGA-3' |
| *Sqstm1* | forward | 5'-TGTGGAACATGGAGGGAAGAG-3' |
| | reverse | 5'-TGTGCCTGTGCTGGAACTTTC-3' |
| *Lc3b* | forward | 5'-TGTGTAACTGTCTCTGTAAG -3' |
| | reverse | 5'-TCTTCTGTTGCTGTTGTC -3' |
| *Ulk1* | forward | 5'- CCCAGCACTACGATGGAAAG -3' |
| | reverse | 5'-CATAAAACAGGCGCAAATCC-3' |
| *Rb1cc1* | forward | 5'-GGAATCTCTGGTCAGGAAGTGC -3' |
| | reverse | 5'-GTCCAAGGCATACAGCCGATCTCCCAGCACTACGATGGAAAG-3' |
